# plastiC: A pipeline for recovery and characterization of plastid genomes from metagenomic datasets

**DOI:** 10.1101/2022.12.23.521586

**Authors:** Ellen S. Cameron, Mark L. Blaxter, Robert D. Finn

**Affiliations:** EMBL-EBI European Bioinformatics Institute, Wellcome Genome Campus, Hinxton, Cambridge CB10 1SD, United Kingdom; Tree of Life, Wellcome Sanger Institute, Wellcome Genome Campus, Hinxton, Cambridge, CB10 1SA, United Kingdom

## Abstract

The use of culture independent molecular methods, often referred to as metagenomics, have revolutionized the ability to explore and characterize microbial communities from diverse environmental sources. Most metagenomic workflows have been developed for identification of prokaryotic and eukaryotic community constituents, but tools for identification of plastid genomes are lacking. The endosymbiotic origin of plastids also poses challenges where plastid metagenomic assembled genomes (MAGs) may be misidentified as low-quality bacterial MAGs. Current tools are limited to classification of contigs as plastid and do not provide further assessment or characterization of plastid MAGs. *plastiC* is a workflow that allows users to identify plastid genomes in metagenome assemblies, assess completeness, and predict taxonomic association from diverse environmental sources. *plastiC* is a Snakemake workflow available at https://github.com/Finn-Lab/plastiC. We demonstrate the utility of this workflow with the successful recover of algal plastid MAGs from publicly available lichen metagenomes.

## 1. Introduction

Culture-independent sequencing of environmental samples, often referred to as metagenomics, has revolutionised the field of microbial ecology, allowing taxonomic and functional characterization of microbial communities from diverse environmental sources. A variety of bioinformatic tools and pipelines are available for the generation and quality estimation of prokaryotic and eukaryotic metagenomic assembled genomes (MAGs) (Parks *et al*., 2015; Saary *et al*., 2020). Organellar genomes are essential to the functioning of eukaryotic organisms, but are poorly assessed in current workflows. Plastid (or chloroplast) genomes are of particular interest because of their roles in primary production, yet the diversity of plastids present in metagenomic data is usually ignored and constrains the full understanding of community structure and function. In standard metagenomic workflows, plastid MAGs may be mistakenly flagged as low-quality bacterial genomes as they lack all of the single copy bacterial marker genes due to the endosymbiotic origin of organelles resulting in potential discard from downstream analyses.

Most existing MAG quality assessment methods are rooted in single-copy marker gene (SCMG) approaches. However, SCMG approaches are not easily transferred to the genomes of plastid organelles due to their reduced genome size, variable gene content (especially driven by organellar-nuclear gene transfer) and large repetitive regions. Thus, alternative methods for quality evaluation are required. Here, we present *plastiC*, a Snakemake (Mölder *et al*., 2021) workflow for the identification and evaluation of plastids in metagenomic samples (www.github.com/Finn-Lab/plastiC) and demonstrate the utility of this workflow to successfully recover algal plastid genomes from lichen metagenomes.

## 2. Methods

### 2.1 Implementation

*plastiC* is designed to analyse contigs generated from metagenomic assembly workflows or high fidelity long-reads (Figure 1). Contigs from these assemblies can be sorted into bins using intrinsic sequence and coverage features, and these bins processed to identify those that putatively correspond to prokaryotic and/or eukaryotic MAGs. These MAGs are typically processed to identify likely taxonomic affiliation derived from marker gene placement, and their quality (*i.e*. completeness and contamination) assessed. While *plastiC* can be run on raw assemblies, we recommend removing contigs that are assigned to “high quality” bins/MAGs before analysis. Putative plastid contigs are identified using *Tiara* (Karlicki *et al*., 2022). As plastid genome are expected to have consistent coverage and tetranucleotide frequency, the assembly is subsequently (re-)binned using *metaBAT2* (Kang *et al*., 2019) with a reduced bin size of 50 kb to ensure that putative plastid bins are retained. The resulting bins are subsequently scanned for plastid contigs as identified by *Tiara*, and bins containing more than a user-specified threshold (default: 90%) of plastid sequences are retained as probable plastid bins.

**Figure 1:**
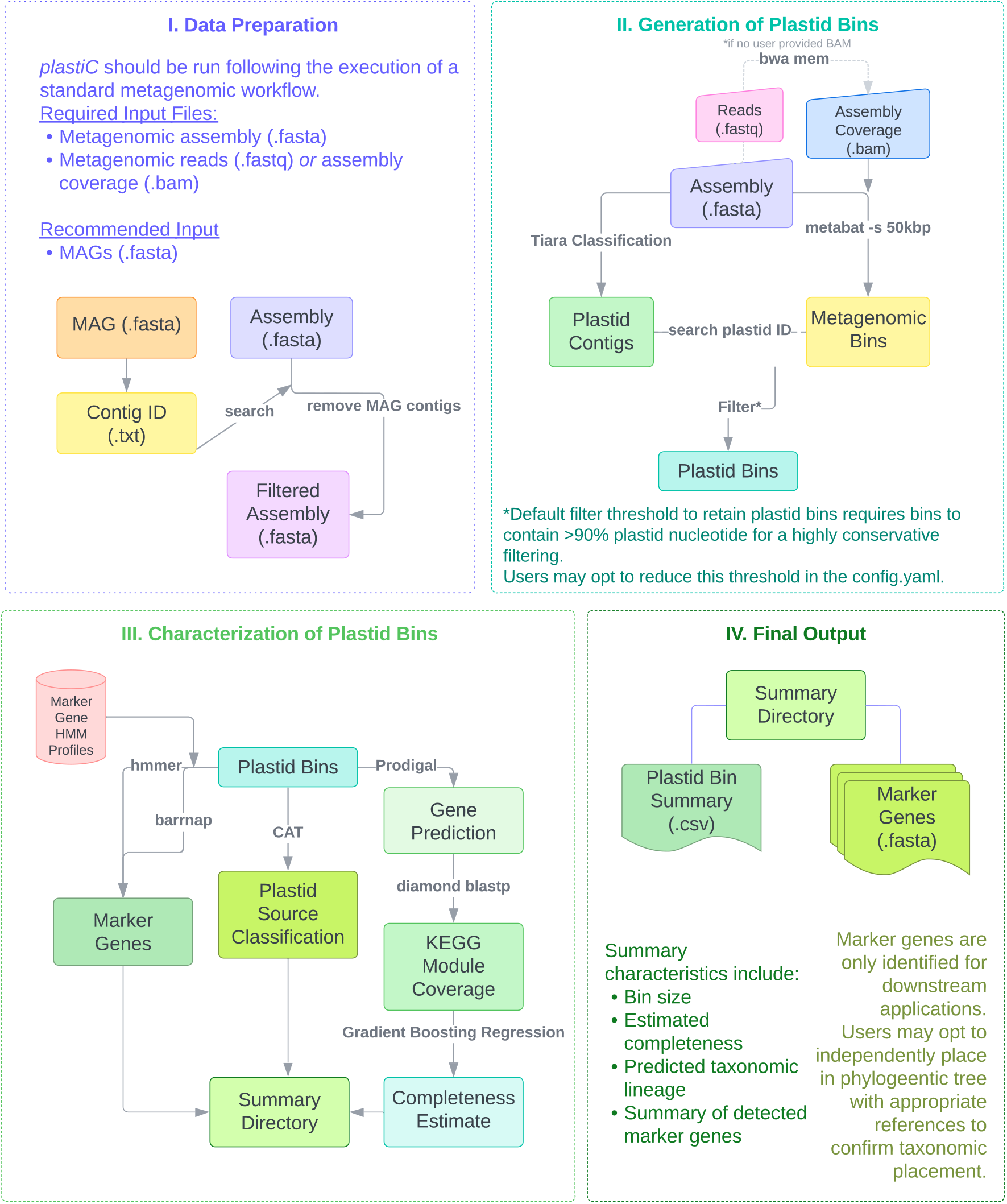
Data preparation and overall workflow for *plastic*. Users provide metagenomic assemblies or high fidelity long reads from which putative plastid bins are identified. A summary report and FASTA files of identified marker loci are generated for each sample.

Machine learning algorithms trained on KEGG module completeness were recently demonstrated as an effective approach for estimating metagenomic bin completeness (Chklovski *et al*., 2022) and we adapted this approach in *plastiC. Prodigal* (Hyatt *et al*., 2010) is used to predict genes in the likely plastid bins, and these predictions are compared to the UniRef100 database (Bateman *et al*., 2021) using *diamond blastp* (Buchfink *et al*., 2021) to obtain KEGG annotations (Kanehisa *et al*., 2016). plastiC then uses the KEGG annotations to calculate KEGG module coverage, and thereby estimate plastid completeness. Due to the presence of shared metabolic pathways and genes between plastid and bacterial chromosomes, bacterial bins may also score as highly complete using this approach, hence the requirement to remove bacterial MAGs before running *plastiC*.

Importantly, *plastiC* also identifies the potential eukaryotic source of the plastid. *plastiC* initially performs a taxonomic classification on the plastid contigs using *CAT* (von Meijenfeldt *et al*., 2019) and additionally searches for taxonomic marker genes using *hmmer* (for *rbcL*) (Eddy, 2022) and *barrnap* (for rRNA loci) (Seeman, 2022). These loci can be used for further taxonomic analyses. As outputs, *plastiC* produces report files summarising bin span, estimated completeness, and taxonomic allocation for each putative plastid bin, as well as FASTA files of the marker loci identified.

### 2.3 Model Training, Testing & Validation

Reference protein sequences from complete plastid genomes available in RefSeq (n = 2633) using Entrez Direct (Kans, 2022) were downloaded on 2022-11-2. One tenth (261) of these references were randomly removed to generate an independent testing set. The training set (n = 2372) was further refined to exclude any reference genomes that were smaller than the median plastid genome size of protists (94 kb) to ensure that outlier small genomes do not impact the training of the model for a final training set of 2279 plastid genomes. *Diamond blastp* was run on both training and testing datasets with UniRef100 to obtain KEGG annotations, which were used to calculate KEGG module completeness. For training sets, reference genome KEGG annotation counts were subsampled without replacement to create simulated examples of plastid genomes with lower levels of completeness ranging from 0 – 100% in increments of 5%. Test reference genomes were also subsampled to create simulated examples to validate the robustness of the completeness estimates ranging from 10 – 100% in increments of 10%.

*Scikit-learn* was utilised to develop and validate machine learning models for estimating metagenomic plastid genome completeness. Specifically, Ada boosting, gradient boosting and random forest regressions were evaluated to determine the effectiveness for estimating completeness of plastid genomes. Training data were split 90 training set:10 test set. K-folds (n=5) cross-validation with shuffling was performed to cross-validate the model.

Each model was tested on reference plastid genomes that were not used in the training set. In addition to the test plastid genome set, KEGG module completeness was predicted for a set of mitochondrial genomes (n = 142) to evaluate whether the model could accurately differentiate between different organellar genomes. Across the cross-validated model set, all three regression models were able to differentiate between plastid and mitochondrial completeness (*i.e*., predict low completeness for mitochondria; high completeness for plastids; Table 1). However, the Ada boosting regression had the lowest plastid completeness estimate and highest mitochondrial estimate of tested models. In combination with the higher mean-squared error, this suggests that the Ada boosting regression model is not the best performing model for application in plastid completeness estimates.

**Table 1:**
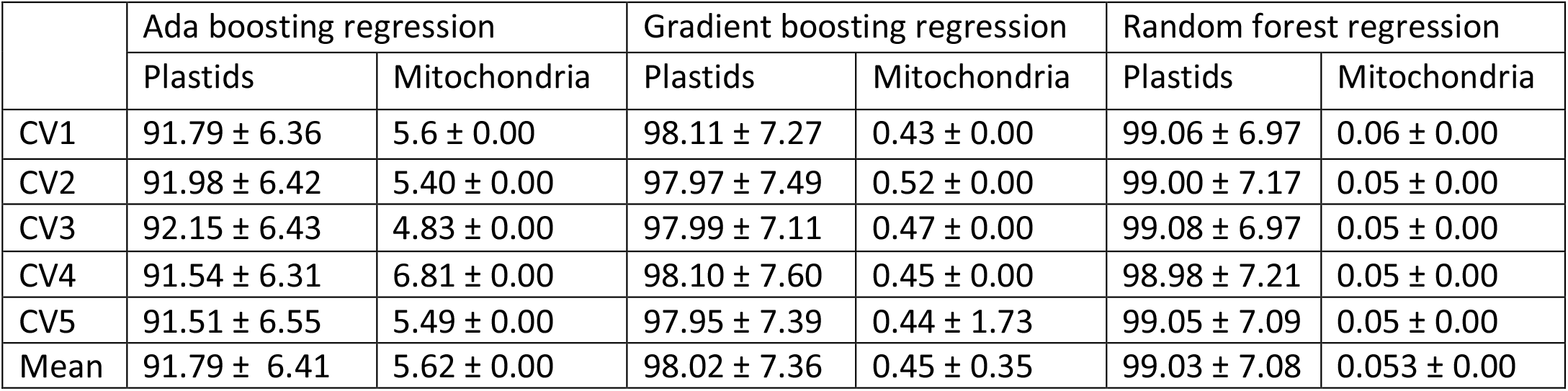
Median predicted completeness values ± standard deviation on whole plastid and mitochondrial reference genomes with k-fold cross validation (n = 5; shuffled) to ensure differentiation between organellar genome completeness scores.

|  | Ada boosting regression |  | Gradient boosting regression |  | Random forest regression |  |
| --- | --- | --- | --- | --- | --- | --- |
|  | Plastids | Mitochondria | Plastids | Mitochondria | Plastids | Mitochondria |
| CV1 | 91.79 ± 6.36 | 5.6 ± 0.00 | 98.11 ± 7.27 | 0.43 ± 0.00 | 99.06 ± 6.97 | 0.06 ± 0.00 |
| CV2 | 91.98 ± 6.42 | 5.40 ± 0.00 | 97.97 ± 7.49 | 0.52 ± 0.00 | 99.00 ± 7.17 | 0.05 ± 0.00 |
| CV3 | 92.15 ± 6.43 | 4.83 ± 0.00 | 97.99 ± 7.11 | 0.47 ± 0.00 | 99.08 ± 6.97 | 0.05 ± 0.00 |
| CV4 | 91.54 ± 6.31 | 6.81 ± 0.00 | 98.10 ± 7.60 | 0.45 ± 0.00 | 98.98 ± 7.21 | 0.05 ± 0.00 |
| CV5 | 91.51 ± 6.55 | 5.49 ± 0.00 | 97.95 ± 7.39 | 0.44 ± 1.73 | 99.05 ± 7.09 | 0.05 ± 0.00 |
| Mean | 91.79 ± 6.41 | 5.62 ± 0.00 | 98.02 ± 7.36 | 0.45 ± 0.35 | 99.03 ± 7.08 | 0.053 ± 0.00 |

The performance of models with plastids of varying completeness was assessed by subsampling the test genomes to simulate varying levels of completeness. Predicted completeness values were compared to expected values at the subset levels to determine efficacy of the model on accurate estimation. Median prediction values and standard deviation was calculated for each iteration of the model produced through cross-validation (Table 2). A linear regression was performed (Figure 2; Table 3) on the predicted completeness compared to expected value and Pearson’s correlation R^2^ was calculated to identify similarity between predicted and expected completeness scores. All regressions and correlation coefficients were statistically significant (p < 2e-16) but the gradient boosting regression model showed the highest correlation between expected and predicted values.

**Table 2:**
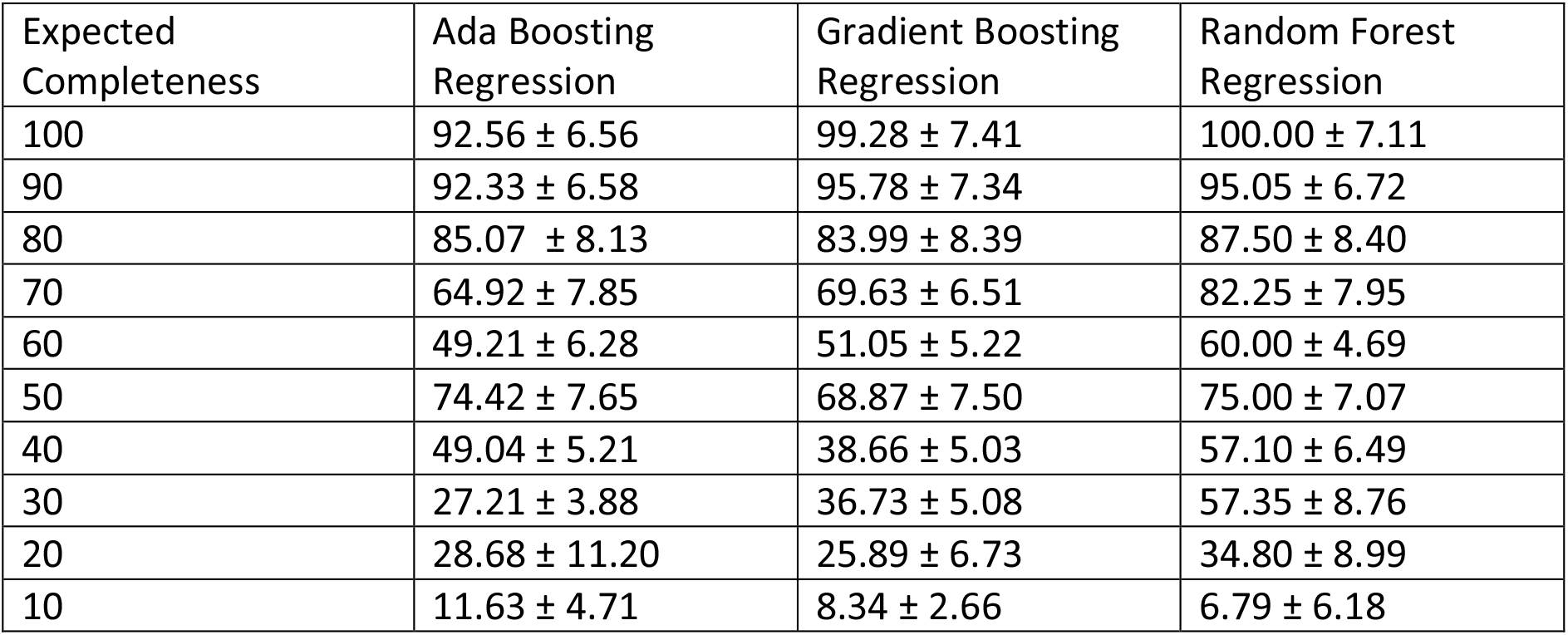
Median predicted completeness values for plastid reference genomes in independent testing validation set (n = 261). Test set plastid references were subsampled down to 10% to evaluate the quality of completeness estimates provided to non-complete genomes.

| Expected Completeness | Ada Boosting Regression | Gradient Boosting Regression | Random Forest Regression |
| --- | --- | --- | --- |
| 100 | 92.56 ± 6.56 | 99.28 ± 7.41 | 100.00 ± 7.11 |
| 90 | 92.33 ± 6.58 | 95.78 ± 7.34 | 95.05 ± 6.72 |
| 80 | 85.07 ± 8.13 | 83.99 ± 8.39 | 87.50 ± 8.40 |
| 70 | 64.92 ± 7.85 | 69.63 ± 6.51 | 82.25 ± 7.95 |
| 60 | 49.21 ± 6.28 | 51.05 ± 5.22 | 60.00 ± 4.69 |
| 50 | 74.42 ± 7.65 | 68.87 ± 7.50 | 75.00 ± 7.07 |
| 40 | 49.04 ± 5.21 | 38.66 ± 5.03 | 57.10 ± 6.49 |
| 30 | 27.21 ± 3.88 | 36.73 ± 5.08 | 57.35 ± 8.76 |
| 20 | 28.68 ± 11.20 | 25.89 ± 6.73 | 34.80 ± 8.99 |
| 10 | 11.63 ± 4.71 | 8.34 ± 2.66 | 6.79 ± 6.18 |

**Figure 2:**
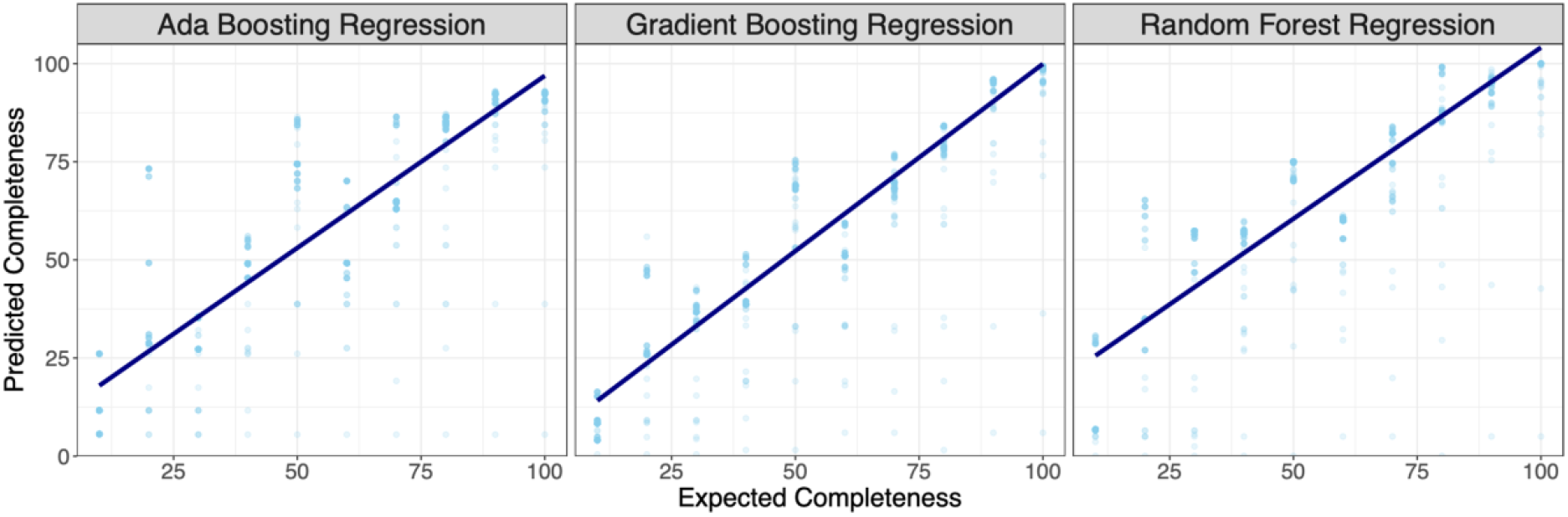
Predicted completeness estimates on test plastid genomes (not used in model training) with three different regression models: Ada boosting, gradient boosting and random forest. The resulting linear equation on linear regression of predicted ∼ expected values is plotted to demonstrate the predictive performance of the different models.

**Table 3:** Linear regression and Pearson correlation as calculated comparing the predicted completeness to expected completeness on the independent test plastid genome estimations.

|  | Ada Boosting Regression | Gradient Boosting Regression | Random Forest Regression |
| --- | --- | --- | --- |
| Linear Equation | $y = 0.88x + 9.19$ | $y = 0.95x + 4.58$ | $y = 0.87x + 16.85$ |
| Adjusted $R^2$ | 0.83 | 0.89 | 0.83 |
| p-value (Linear regression) | $<2.20e-16$ | $<2.20e-16$ | $<2.20e-16$ |
| Pearson's Correlation | 0.91 | 0.95 | 0.91 |
| p-value (Pearson's) | $<2.20e-16$ | $<2.20e-16$ | $<2.20e-16$ |

To conclude evaluation of the effectiveness of each model, differences between the expected and predicted values for each cross-validated model iteration were calculated (Figure 3). The random forest regression model had the best performing mean-squared error, highest completeness estimates for whole plastid genomes and lowest completeness estimates for whole mitochondrial genomes. However, it frequently overestimated completeness (median difference = -7.5). Based on the correlation between expected and predicted and median value of discrepancy between predicted and expected values (n = 0.37), the gradient boosting regression model was chosen as the best-performing model for plastid genome completeness estimates.

**Figure 3:**
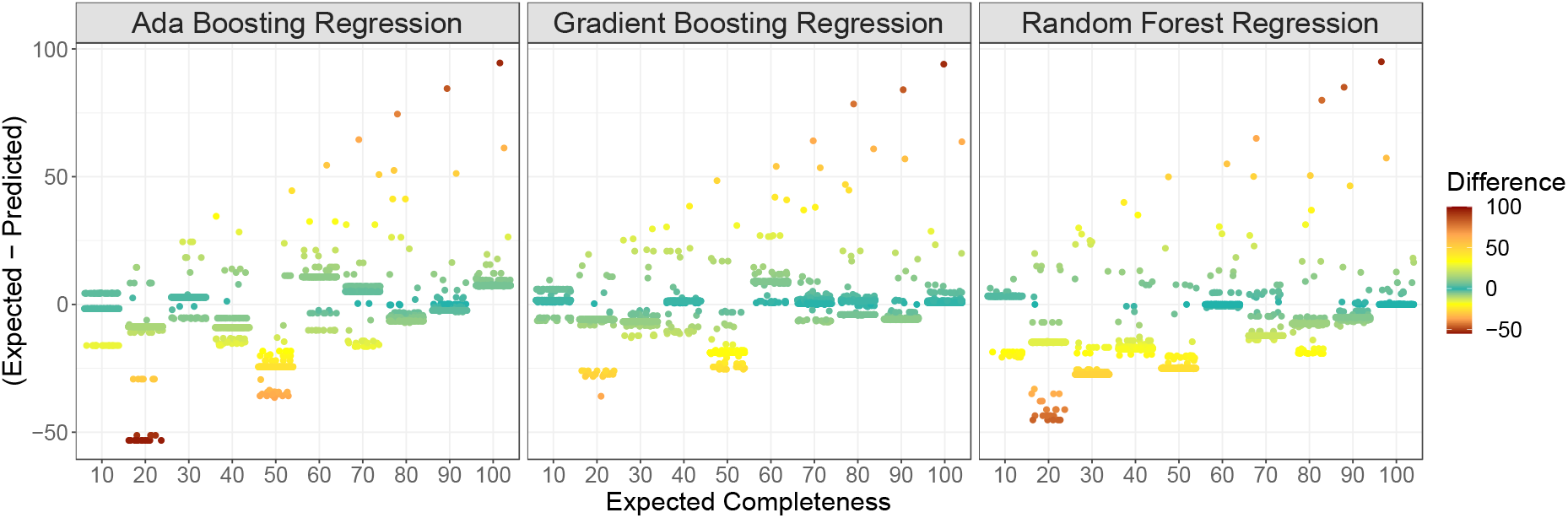
Differences in expected completeness of subsampled plastid reference genomes and predicted completeness. Positive values in difference indicate underestimation of plastid predicted completeness while negative values represent overestimation. Gradient boosting regression consistently had the smallest median discrepancy between predicted values and expected value (0.37), while random forest had the largest discrepancy frequently resulting in the overestimation of predicted plastid completeness (median = -7.5). Ada boosting regression performed moderately (median = -2.12).

### 2.3 Operation

*plastiC* is a Snakemake workflow which automates the process of identification, assessment and taxonomic allocation of plastid genomes in metagenomic samples. Specifically, this workflow contains eight rules, including an optional mapping step. Users are required to populate the *config.yaml* providing information on paths to directories and files, and options for workflow customization (e.g., filter thresholds, inclusion of mapping rule). *plastiC* requires one of the following for each sample to be analysed: i) a metagenomic assembly, ii) high fidelity long-reads. If available, users also have the option to provide a *bam* file for mapped reads to the assembly. Users may opt to decrease the minimum percentage of plastid sequence required to be classified as a probable plastid bin (*min_plastid_content*) to increase or decrease stringency as appropriate for individual use cases.

Since *plastiC* runs on previously generated metagenomic assemblies, memory requirements are typically low but may vary depend on the nature of the sample. In development, the majority of rules required less than 1 GB RAM for successful completion with the exception of i) contig classification using *Tiara* (<5 GB RAM) and ii) taxonomic source classification using *CAT* (<8 GB RAM). These memory requirements may vary depending on sample size and complexity of the community. Users may specify computational resource allocations for individual rules and execution on a high-performance compute cluster in the *cluster.yaml*.

## 3. Use Cases

To demonstrate the effectiveness of *plastiC* on metagenomic data, lichen metagenomes were downloaded from the project accession PRJNA646656 (n = 13; Smith et al., 2020) in the European Nucleotide Archive. Lichens are composite organisms composed of a mutualist symbiotic association between a primary fungal partner (mycobiont) and algal partner (photobiont). Chlorophyta (green alga) taxa are the photobionts in many lichen species and thus plastids should be present in lichen metagenomic samples. The analysed samples were derived from 10 species of lichen spanning 6 genera (Table 5) which are all expected to contain *Trebouxia*, a green algal genus, as their primary photobiont. Metagenomic assemblies for these samples were generated as part of a large survey of lichen microbial symbionts (Tagirdzhanova et al., 2023). In brief, metagenomic datasets were filtered using *fastp* and human contamination was removed using *BMTagger*. Quality-controlled reads were assembled using *metaSPAdes*. These assemblies were used with *plastiC* to recover plastid genomes.

Plastid contigs were identified in all 13 samples using *Tiara* (Table 4). Metagenomic assemblies were binned using *metaBAT2* with the reduced bin size threshold of 50 kb. These bins were then classified as being putative plastid genomes based on the fraction of plastid contigs. Bins that were comprised of >90% plastid nucleotides were retained as probable plastid bins for further analysis. Of the 13 lichen metagenomes analysed, a single plastid bin was identified in 8 of them (Table 4). For the remaining 5 samples, plastid contigs were not successfully binned and were retained in the unbinned portion with other sequences.

**Table 4:** Lichen metagenome assembly and binning information. Metagenomes were assembled into contigs and these assemblies were used to identify plastid contigs using *Tiara* and for binning with *metaBAT2*. Probable plastid bins were identified based on the distribution and location of plastid contigs within bins, with a threshold of >90% to be retained for downstream analyses.

| Run Accession | Scientific Name | Total Contigs | Plastid Contigs | Total Bins | Probable Plastid Bins |
| --- | --- | --- | --- | --- | --- |
| SRR12240187 | *Xanthoparmelia chlorchroa* | 190210 | 12 | 15 | 1 |
| SRR12240188 | *Xanthoparmelia chlorochroa* | 115748 | 12 | 18 | 1 |
| SRR12240177 | *Xanthoparmelia maricopensis* | 364371 | 20 | 23 | 1 |
| SRR12240174 | *Xanthoparmelia neocumberlandia* | 253456 | 108 | 23 | 0 |
| SRR12240175 | *Xanthoparmelia neocumberlandia* | 171912 | 93 | 13 | 1 |
| SRR12240180 | *Xanthoparmelia chlorochroa* | 164812 | 14 | 20 | 1 |
| SRR12240179 | *Xanthoparmelia mexicana* | 253280 | 3 | 24 | 1 |
| SRR12240178 | *Xanthoparmelia plittii* | 190261 | 81 | 17 | 1 |
| SRR12240185 | *Mobergia calculiformis* | 129614 | 505 | 28 | 1 |
| SRR12240183 | *Physcia biziana* | 330121 | 64 | 27 | 0 |
| SRR12240182 | *Physciella chloantha* | 650113 | 31 | 26 | 0 |
| SRR12240181 | *Rinodina sp* | 170759 | 207 | 28 | 0 |
| SRR12240184 | *Oxernella safavidorum* | 455310 | 587 | 22 | 0 |

**Table 5:** Probable plastid bin characteristics including bin span, total contig count and number of plastid contigs. Completeness estimates were performed based on KEGG module coverage and gradient boosting regression, and taxonomic association of plastid genomes performed with *CAT*.

| Run Accession | Bin Span | Total Contig Count | Number of Plastid Contigs | Completeness Estimate | Taxonomic Association |
| --- | --- | --- | --- | --- | --- |
| SRR12240187 | 231882 | 1 | 1 | 96.1 | *Trebouxia* |
| SRR12240188 | 231734 | 2 | 2 | 96.1 | *Trebouxia* |
| SRR12240177 | 238137 | 10 | 10 | 95.9 | *Trebouxia* |
| SRR12240174 | N/A | N/A | N/A | N/A | N/A |
| SRR12240175 | 121980 | 15 | 14 | 33.4 | *Trebouxia* |
| SRR12240180 | 254207 | 5 | 5 | 96.1 | *Trebouxia* |
| SRR12240179 | 258520 | 1 | 1 | 96.4 | *Trebouxia* |
| SRR12240178 | 64547 | 11 | 10 | 10.8 | *Trebouxia* |
| SRR12240185 | 77893 | 9 | 8 | 19.1 | *Trebouxia* |
| SRR12240183 | N/A | N/A | N/A | N/A | N/A |
| SRR12240182 | N/A | N/A | N/A | N/A | N/A |
| SRR12240181 | N/A | N/A | N/A | N/A | N/A |
| SRR12240184 | N/A | N/A | N/A | N/A | N/A |

Taxonomic source prediction was performed on the putative plastid bins. All plastid bins identified in the sample were attributed to the genus *Trebouxia*, which corresponds with expectations of the photobiont in these lichens being a trebouxoid green alga. Plastid bins ranged in estimated completeness from 10.8 to 96.4% and completeness was positively correlated to the bin span in these example samples (Figure 4).

**Figure 4:**
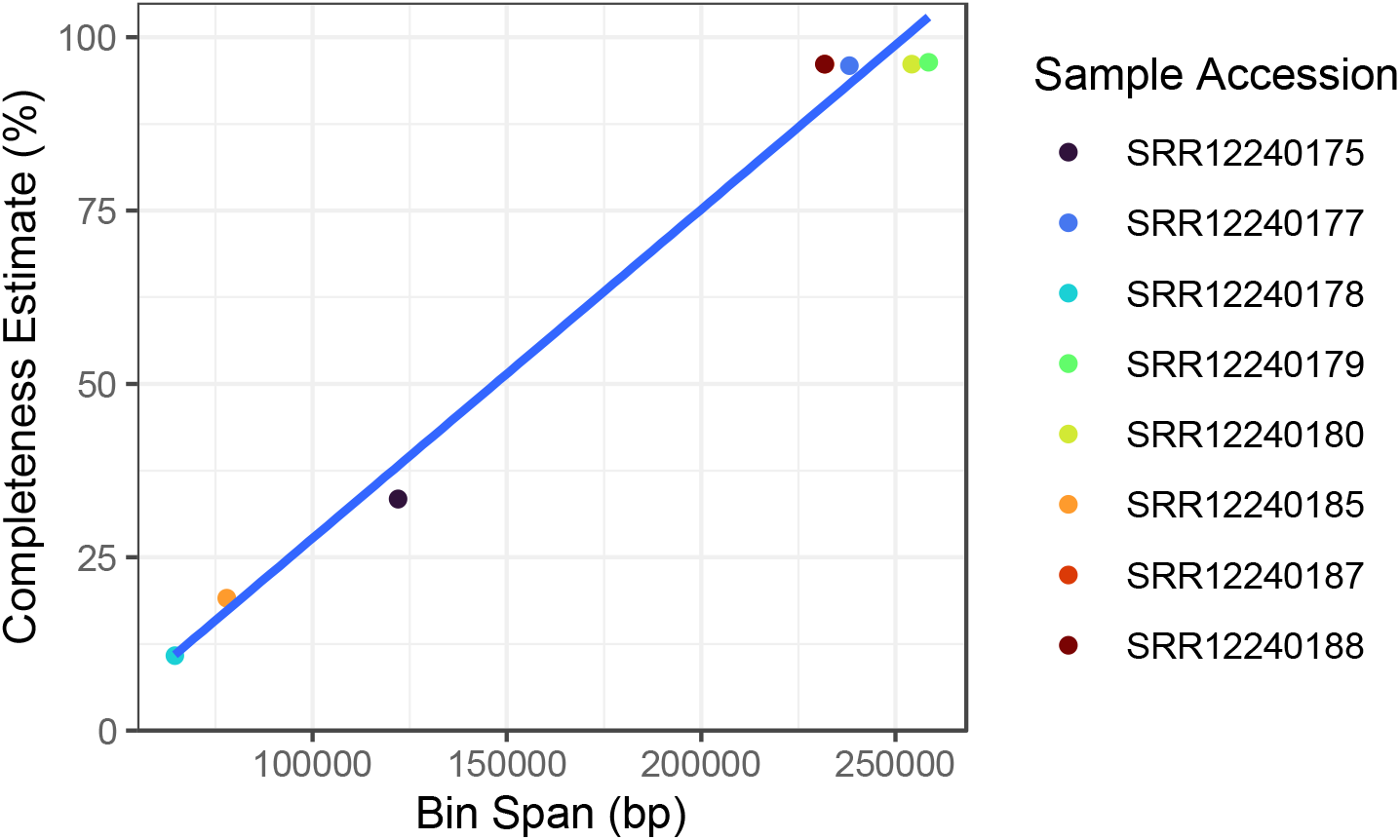
Recovered *Trebouxia* plastid MAGs from lichen metagenomes. Samples with lower completeness plastid MAGs correlate with smaller bin sizes suggesting incomplete plastid genome content present in metagenomic data.

## 4. Discussion

Currently, metagenomic characterization of plastid genomes is limited to the identification of plastid contigs using *Tiara*. However, only identifying plastid contigs prevents full disentanglement of metagenomic datasets, including the potential presence of multiple plastid genomes arising from different sources. *plastiC* uniquely provides users with the opportunity to characterize plastid genomes including separation into independent plastid genomes and source classification to inform on present eukaryotic photosynthetic organisms. Notably, this is the first tool to provide a completeness estimate for plastid genomes from metagenomic sources. The high copy number and small genome size of plastids may permit easier recovery and serve as a proxy for the identification of the presence of eukaryotes which may not have sufficient genomic signal to be recovered as a high-quality eukaryotic MAG, and facilitate targeted coassembly of samples, based on common plastid composition. Furthermore, plastiC offers the potential to expand reference databases with metagenomic derived plastids. *plastiC* will support characterization of plastids in diverse ecological contexts including, for example, symbiotic interactions (e.g., lichens as demonstrated in the Use Cases), complex marine ecosystems and characterisation of components of herbivore diets.

## 5. Acknowledgements

This research was supported by EMBL (European Molecular Biology Laboratory) core funds. We would also like to acknowledge Paul Saary for discussions on metagenomic quality estimation and Richard Challis for advice on the production of user-friendly Snakemake workflows.

## Notes

https://github.com/finn-lab/plastic

### Competing Interest Statement

The authors have declared no competing interest.

### Summary of Updates

Including supplementary information on model training and use cases in the main body of the text. Addition of Figure 4 to visualise completeness estimate correlation to plastid genome size in lichen use case.

